# *Salmonella* Typhi acquires diverse plasmids from other Enterobacteriaceae to develop cephalosporin resistance

**DOI:** 10.1101/2020.10.16.343467

**Authors:** Jobin John Jacob, Agila K Pragasam, Karthick Vasudevan, Balaji Veeraraghavan, Gagandeep Kang, Jacob John, Vasant Nagvekar, Ankur Mutreja

## Abstract

**Background:** Recent reports have established the emergence and dissemination of extensively drug resistant (XDR) H58 *Salmonella* Typhi clone in Pakistan. In India where typhoid fever is endemic, only sporadic cases of ceftriaxone resistant *S*. Typhi are reported. This study aimed at elucidating the phylogenetic evolutionary framework of ceftriaxone resistant *S*. Typhi isolates from India to predict their potential dissemination in endemic regions.

**Methods:** Five ceftriaxone resistant *S*. Typhi isolates from three tertiary care hospitals in India were sequenced on an Ion Torrent Personal Genome Machine (PGM). A core genome single-nucleotide-polymorphism (SNP) based phylogeny of the isolates in comparison to the global collection of MDR and XDR *S*. Typhi isolates was built. Two of five isolates were additionally sequenced using Oxford Nanopore MinION to completely characterize the plasmid and understand its transmission dynamics within Enterobacteriaceae.

**Results:** Comparative genomic analysis and detailed plasmid characterization indicate that while in Pakistan (4.3.1 lineage I) the XDR trait is associated with *bla*_CTX-M-15_ gene on IncY plasmid, in India (4.3.1 lineage II), the ceftriaxone resistance is due to short term adaptation of resistance plasmids such as IncX3 or IncN.

**Conclusion:** Since the bacterial acquisition of smaller resistance plasmids such as IncX3 or IncN from other Enterobacteriaceae can be much faster than the larger IncY plasmids, the rapid expansion of these genotypically novel XDR *S*. Typhi could potentially cause large outbreaks. Therefore, continuous monitoring of *S*. Typhi lineages carrying cephalosporin resistance on IncX3 or IncN plasmids is vital not just for India but globally.

**Importance:** Genomic analysis of cephalosporin resistant *S*. Typhi isolated from India indicates the potential of *S*. Typhi to develop cephalosporin resistance by acquiring diverse plasmids from other Enterobacteriaceae. We identified the occurrence of independent acquisition of drug-resistant plasmids such as IncX3 and IncN with genes encoding beta-lactamases in H58/4.3.1.2 lineage. A short term adaptation of drug-resistant plasmids in H58/4.3.1.2 lineage can be the reason for the sporadic cases cephalosporin resistant *S*. Typhi in India. However, the IncY plasmid acquired by isolates that belong to H58/4.3.1.1 lineage appeared to be well adapted as observed in XDR S. Typhi outbreak in Pakistan. Plasmid acquisition and maintenance of cephalosporin resistant *S*. Typhi appears to be specific to the phylogenetic lineage as lineages differ in compensating the initial cost imposed by the plasmid. The stable maintenance of these resistance plasmids without a fitness cost, are determinant in understanding the future spread of cephalosporin resistance in *S*. Typhi. Therefore, critical strategies in monitoring and control of cephalosporin resistant *S*. Typhi is needed to tackle further public health crisis.

## Introduction

Enteric fever is a severe systemic infection caused primarily by *Salmonella enterica* serovar Typhi and, serovar Paratyphi A [1]. These typhoidal serovars are restricted to human hosts and typically occur in low and middle – income countries (LMIC) [2]. Although direct faecal-oral route is the predominant mode of transmission, recent reports suggest that indirect transmission may also occur as the bacteria can survive for extended periods in the environment [3]. The global burden of typhoid fever is estimated to be between 11 and 21 million with 128, 000 to 161, 000 deaths annually (WHO, 2019). Global data suggests that the majority of the reported enteric fever morbidity and mortality takes place in endemic regions of South Asian, Southeast Asian and African countries [4].

The management of enteric fever is challenging due to the emergence of antibiotic resistant *S*. Typhi strains and their changing resistance profiles [5]. The indiscriminate use of first line antimicrobial agents (ampicillin, chloramphenicol and co-trimoxazole) during the 1960s led to the emergence of multi drug resistance (MDR) in sporadic cases initially, followed by larger outbreaks during 1970-1990 [6]. Fluoroquinolones (FQs), such as ciprofloxacin, then became the drug of choice for treatment of MDR *S*. Typhi. However, the decreased ciprofloxacin susceptibility (DCS) phenotype became dominant globally within a few years, resulting in clinical failures [7]. Currently, ceftriaxone and azithromycin are the drugs of choice. However, there are increasing reports of ceftriaxone resistant and azithromycin resistant *S*. Typhi [8].

*S*. Typhi isolates with extensive drug resistance (XDR) have emerged in Sindh, Pakistan, with resistance to ampicillin, chloramphenicol, co-trimoxazole, fluoroquinolones and third-generation cephalosporins [9]. This large scale outbreak reported a total of 5274 XDR *S*. Typhi cases between 2016 and 2018 (WHO, 2018). The XDR *S*. Typhi isolates carried an IncY plasmid harboring a *bla*_CTX-M-15_ and *qnrS1* gene while the antimicrobial resistance (AMR) cassette conferring resistance to first-line drugs was integrated into the chromosome as a composite transposon [9, 10]. In India where typhoid fever is endemic, only sporadic cases of ceftriaxone resistant *S*. Typhi have been reported [11–13].

Until recently, phylogenetic inferences of the evolution of cephalosporin resistant *S*. Typhi were limited to the reported XDR *S*. Typhioutbreak in Pakistan [9], possibly because only sporadic cases are reported across other locations in South Asia. Since *S*. Typhi H58 lineage can acquire MDR plasmids (p60006) from other Enterobacteriaceae, this event could also occur in other regions of Asia where typhoid is endemic [9]. Our study investigated the evolutionary trajectories of cephalosporin resistancein *S*. Typhi with special reference to the endemic regions in South Asia. We also investigated the role of plasmid in this clonal expansion to predict the possibility of the rise and spread of cephalosporin resistant *S*. Typhi in India.

## Materials and Methods

### Bacterial Isolates, Identification and AST

Five clinical isolates of ceftriaxone resistant *Salmonella* Typhi from three different tertiary care hospitals in India between 2015 and 2018 were confirmed by serotyping according to the *Kauffmann-White* scheme [14] and standard microbiological techniques. Antimicrobial susceptibility testing (AST) was performed by using agar disk diffusion (DD) method for six antimicrobial agents including ampicillin (10 μg), chloramphenicol (30 μg), co-trimoxazole (1.25/23.75 μg), ciprofloxacin (5 μg), ceftriaxone (30 μg) and azithromycin (5 μg). Minimum inhibitory concentration (MIC) for ceftriaxone was determined using the broth micro dilution (BMD) method in accordance with the Clinical and Laboratory Standards Institute (CLSI) 2018 guidelines and interpretative criteria [15].

### DNA Extraction and Whole genome sequencing

Genomic DNA of the study isolates was extracted from an overnight culture (14 - 16 hrs) grown at 37°C on blood agar, using the QIAamp DNA Mini Kit (Qiagen, Hilden, Germany) according to the manufacturer’s protocol. The extracted DNA was subjected to whole genome sequencing (WGS) using the Ion Torrent PGM sequencer (Life Technologies, Carlsbad, CA) with 400 bp read chemistry. Two of five isolates were sequenced using the Oxford Nanopore MinION sequencer (ONT, Oxford, UK) per standard protocol to fully resolve the plasmid structure.

### Genome assembly, Genotyping and Plasmid typing

Hybrid genome assembly was carried out according to the standardized protocol developed in-house [16] using Unicycler hybrid assembly pipeline (v0.4.6). Ion torrent reads assembly was achieved *de novo* in the SPAdes assembler (v 5.0.0.0) embedded in the Torrent suite server (v.5.0.3). The quality metrics of the assembled genome was analysed using Quast(v.4.5) and genomes were annotated using Prokaryotic genome annotation pipeline (PGAP v.4.9) before being submitted to NCBI.

In-*silico* Multi-Locus Sequence Typing (MLST) was determined using Enterobase database (https://enterobase.warwick.ac.uk) available in pubMLST. The resistance profile of the assembled genomes was identified using ResFinderv.3.1 available from (https://cge.cbs.dtu.dk/services/ResFinder/). Isolates were genotyped using the genotyping scheme as described on the GitHub repository (https://github.com/katholt/genotyphi). Plasmid typing was carried out using Plasmid finder available from the CGE server (https://cge.cbs.dtu.dk/services/).

### SNP calling and Phylogenetic tree construction

Core genome single-nucleotide polymorphisms (SNPs) were identified using Snippy v.0.2.6 (https://github.com/tseemann/snippy) with CT18 (NC_003198) as the reference [17]. The recombination within the alignment file was filtered and removed using the Gubbins algorithm v. 2.0.0 [18] and the non-recombinant SNPs, were used to construct the phylogenetic tree using Fast tree. The maximum - likelihood tree with 100 bootstrap values was rooted to the reference genome CT18 and labelled using the interactive tree of life software iTOL v.3 [19].

### Plasmid characterization and comparative genomics

Plasmidsfrom two representative isolates (One isolate each from Mumbai and Vellore) were circularized by combining the reads of Ion torrent and MinIon sequencing platforms as described earlier. The remaining three IncX3 plasmids (carried by Mumbai isolates) were extracted from short reads using plasmidSPAdes [20] with representative plasmids as the reference. The plasmid comparison was visualized and analyzed using CGview server v.1.0 [21].

## Results

### Identification of ceftriaxone resistant *S*. Typhi

All the study isolates werephenotypically and biochemically identified as *S*. Typhi and serologically confirmed by traditional serotyping.The antimicrobial susceptibility profiling of our study isolates showed resistance to ampicillin, ceftriaxone and ciprofloxacin whilst being sensitive to chloramphenicol, co-trimoxazole and azithromycin. Among the five isolates, none were MDR and all showed similar AST patterns. The demographic details and the resistance profiles of the study isolates are given in Table 1.

**Table 1:**
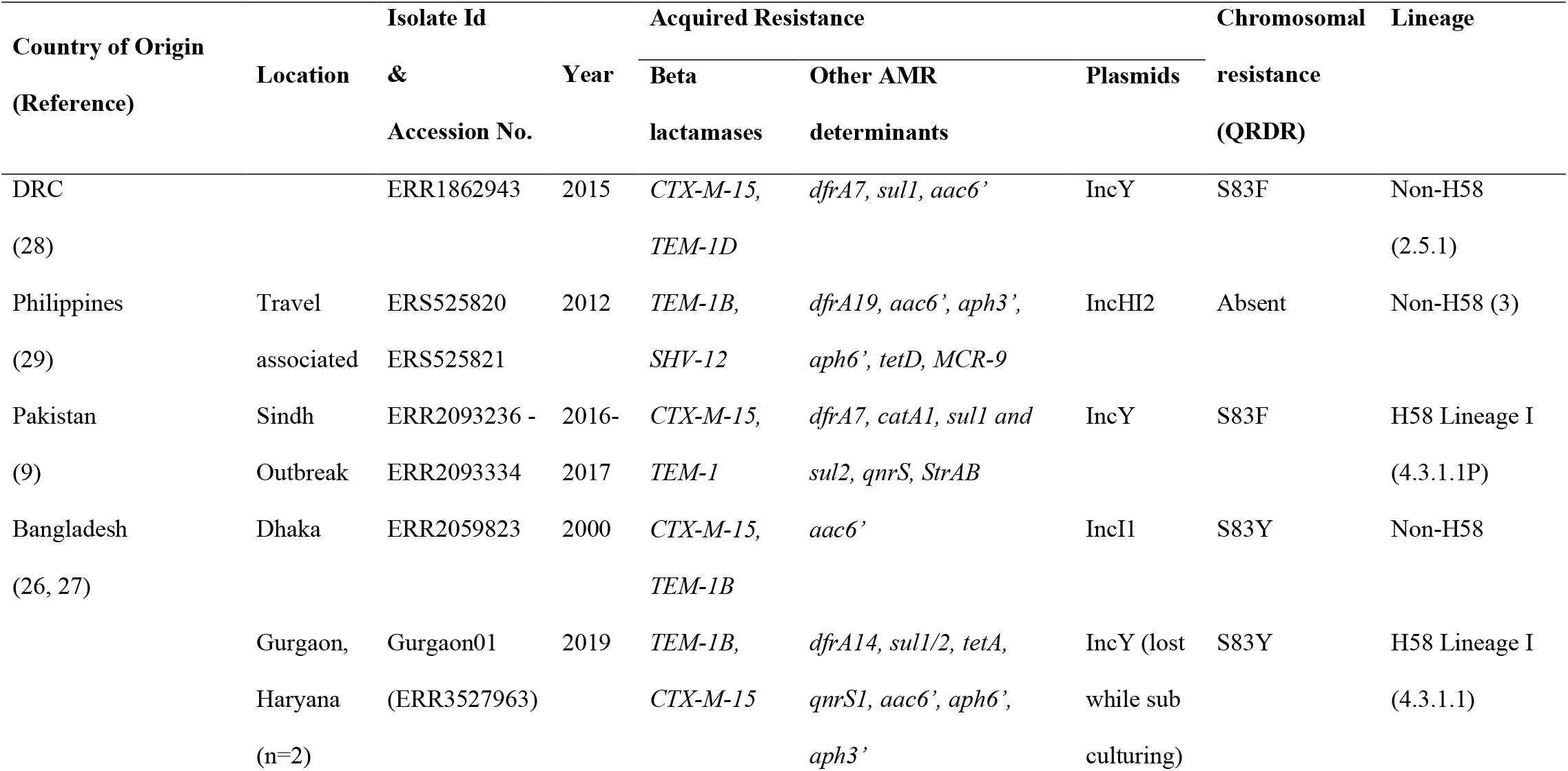
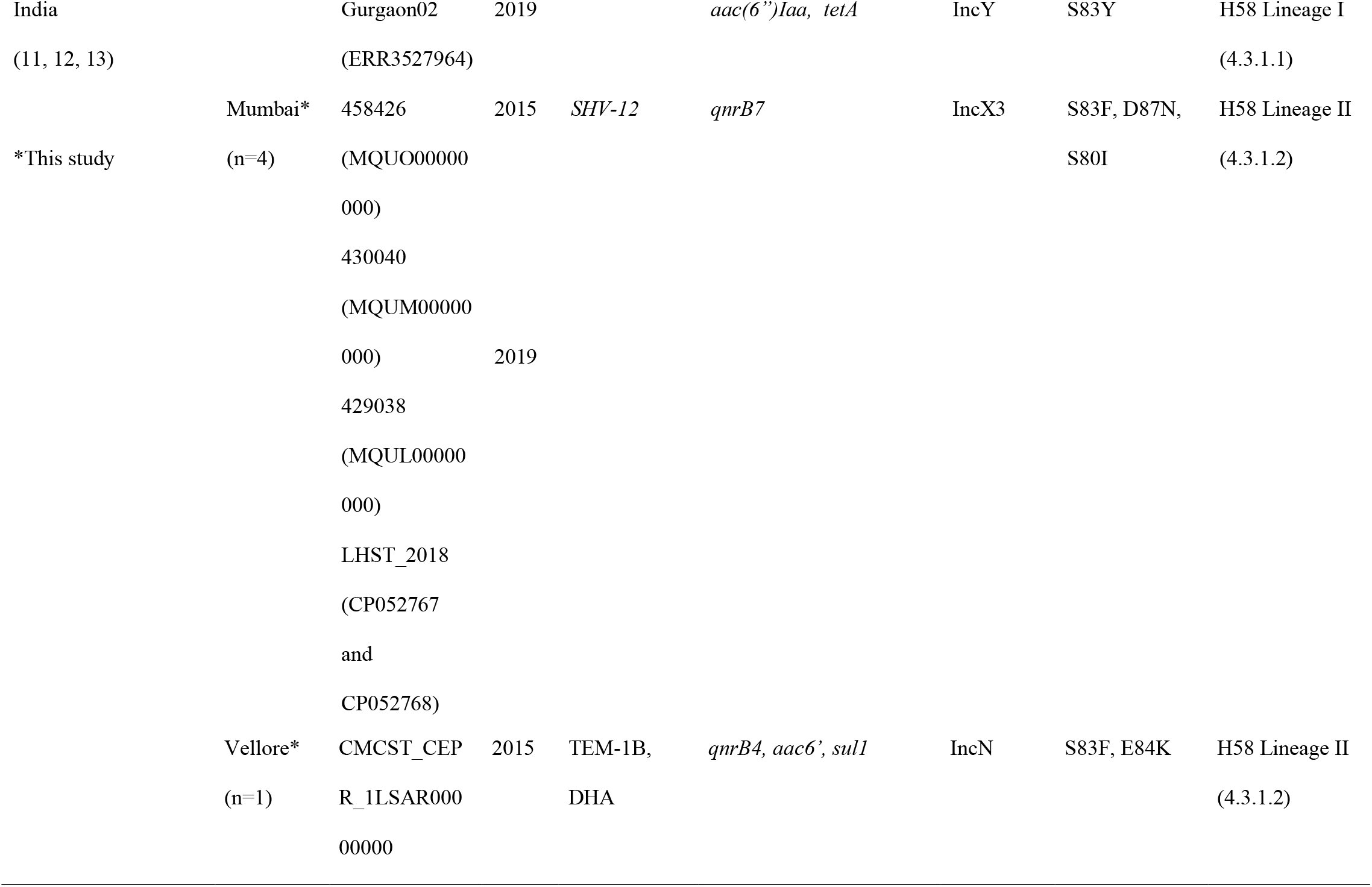
Molecular fingerprint of cephalosporin resistant *S*. Typhi

Genome based MLST analysis classified the isolates to ST1 and identified them as 4.3.1.2 (H58) genotype. Resistome analysis from the whole genome revealed the presence of *bla*_SHV-12_, and *qnrB7* genes in four Mumbai isolates, while the Vellore isolate carried *bla*_*T*EM-1B_, *bla*_DHA-1_, *qnrB4*, aac(6)Iaa, and *sul-1* resistance genes. Ciprofloxacin resistance could be attributed to QRDR triple mutations (gyrA: S83F, D87N, parC: S80I) for the Mumbai isolates and double mutations (gyrA: S83F, parC: E84K) for the Vellore isolate.

### Multiple, independently evolving, ceftriaxone resistant lineages circulating in India

To determine the emergence of ceftriaxone resistance in Indian *S*. Typhi isolates, the phylogenetic relationship of study isolates were compared with the representative isolates (H58 and non-H58) from the global *S*. Typhi collection (**Fig. 1**). Ceftriaxone resistant isolates from the recently reported XDR *S*. Typhi outbreak in the Sindh, Pakistan as well as sporadic ceftriaxone resistant isolates from Gurgaon, India and Dhaka, Bangladesh were included in the phylogenetic tree (Supplementary Table S1). The XDR *S*. Typhi isolates from Pakistan formed a distinct clade (4.3.1.1P) within the H58 lineage. Isolates reported to have originated from a direct travel history from Pakistan were also found to be in the same cluster. The two documented ceftriaxone resistant isolates from Bangladesh clustered with the non-H58 lineage along with the ceftriaxone sensitive isolates from the same location. Interestingly the five study isolates from India did not cluster together, and were distributed across the 4.3.1 lineage II. The four cephalosporin resistant isolates from Mumbai, India formed a subclade within the H58 lineage II and differed by 2 SNPs from the closest isolates within the subclade. However the recently reported ceftriaxone resistant *S*. Typhi from Gurgaon, India clustered within the subclade of 4.3.1 lineage I with a difference of 3 SNPs from the ceftriaxone sensitive isolates. It is therefore likely that all the ceftriaxone resistant *S*. Typhi isolates from India have evolved independently from respective geographical locations.

**Fig. 1:**
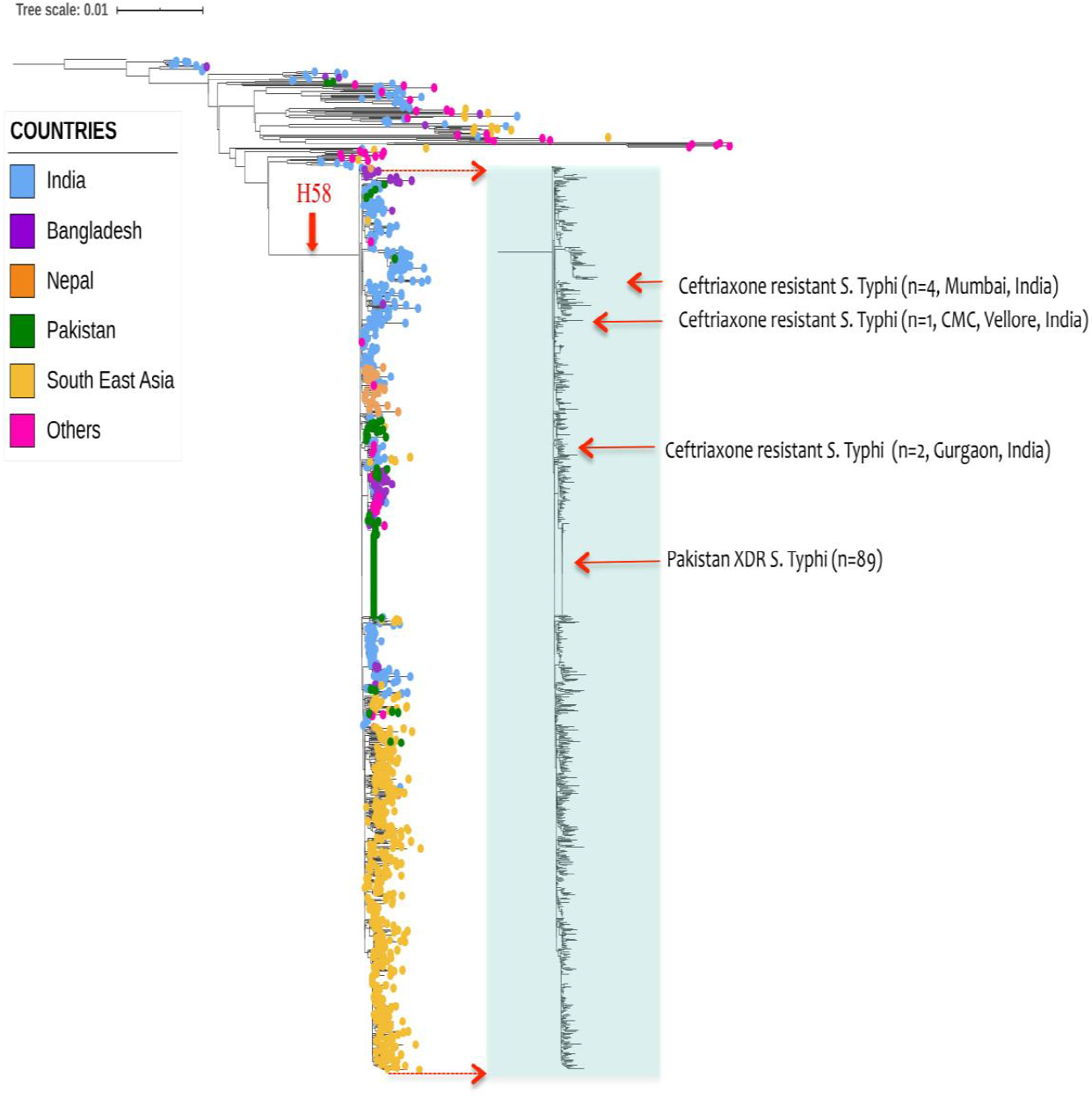
Maximum likelihood tree of 1005 *S*. Typhi (H58 and non-H58) inferred from 2072 SNPs and rooted against the reference genome CT18. Light shaded box indicates non-H58 population, while the rest are H58 group. The tips of the tree are coloured according to the geographical origin of the genomes. Red colored squared boxes with arrows indicates the position of ceftriaxone resistant *S*. Typhi in the context of global phylogeny, indicating similarity with respect to the geographical locations. Tree scale indicates the number of substitutions per genome.

### Non-IncY plasmids carrying ceftriaxone resistance

The complete circular plasmids of two representative isolates were used as a reference to reconstruct plasmids from the short read assembly in other isolates. Four study isolates from Mumbai carried the *bla_SHV-12_* and *qnrB7* AMR genes on an IncX3 plasmid (**Fig. 2**). The antibiotic resistance loci from the plasmid were found to be a composite transposon inserted into the IncX3 backbone. However, the plasmid IncN from the Vellore isolate carried *bla*_TEM-1B_, *bla*_DHA-1_, *qnrB4*, and *sul-1* resistance genes. The IncX3 plasmid responsible for the cephalosporin resistance in *S*. Typhi in India is very closely related to the IncX3 plasmid in other Enterobacteriaceae.

**Fig. 2:**
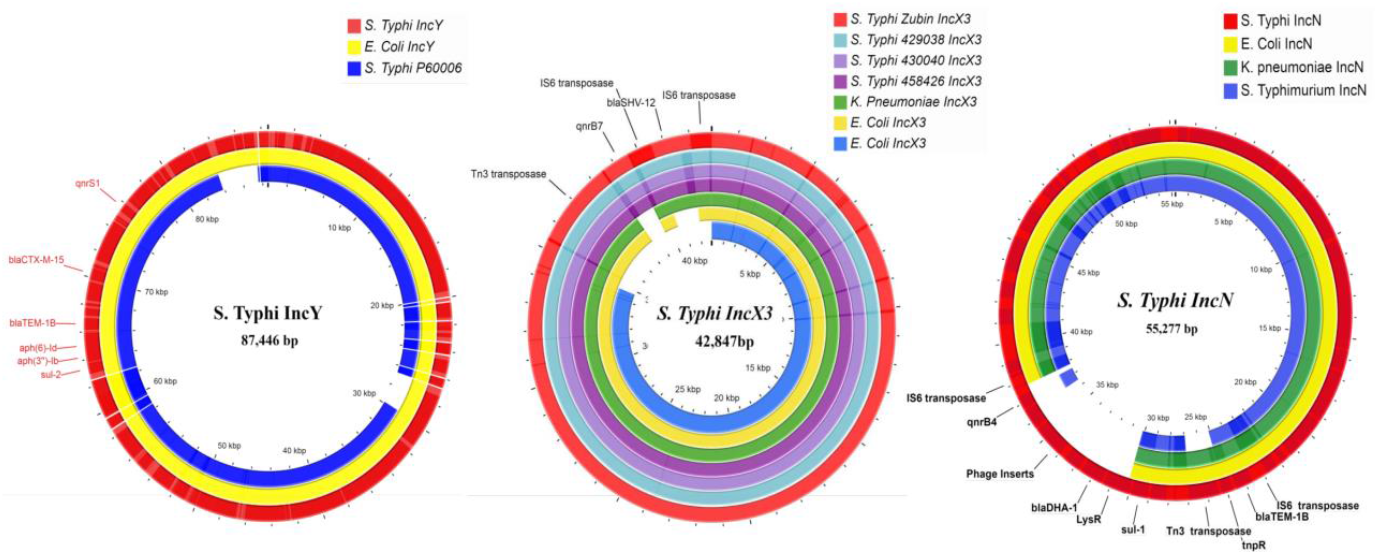
Circular representation of plasmids assembled from *S*. Typhi isolates displayed using CG view server with reference genomes. (a) The IncY plasmid from cephalosporin resistant *S*. Typhi isolate from Gurgaon, India is compared with similar plasmids identified from XDR Pakistan *S*. Typhi (Plasmid p60006) and highly similar *E. coli* IncY plasmid. (b) Comparison of IncX3 plasmid from cephalosporin resistant Indian *S*. Typhi isolates (n=4) with global Enterobacteriaceae associated IncX3 plasmids. (c) The IncN plasmid assembled from cephalosporin resistant *S*. Typhi Vellore isolate was compared with representative isolates from Genbank

## Discussion

Currently, ceftriaxone and azithromycin are the drugs of choice and resistance to these agents challenges the treatment of typhoid fever. This has been demonstrated by the emergence of ceftriaxone resistant *S*. Typhi in Pakistan, with resistance to five classes of antimicrobials, including all three first-line agents, fluoroquinolones and ceftriaxone. This study provides evolutionary insights into the emergence of cephalosporin resistant *S*. Typhi in India to predict possible further clonal expansion.

The emergence of Indian cephalosporin resistant *S*. Typhi appears to be by multiple independent genetic events by acquisition of different resistance plasmids. This is in line with the global emergence of cephalosporin resistance in *S*. Typhi as summarized in Table 1. So far, 18 studies have reported cephalosporin resistant *S*. Typhi across the globe, with the earliest report of *bla*_CTX-M-15_ gene in Western Asia during 2003-2006 [22]. While the majority of reports were based on phenotypic data, a few have reported genetic AMR and plasmid profiles. Except for the outbreak event reported from Pakistan [9], all other reports are of sporadic cases from different locations. Hence there is a lack of a global perspective of the evolution and spread of cephalosporin resistant *S*. Typhi.

In our study, the genetic characterization of cephalosporin resistant *S*. Typhi isolates from India showed that as in Pakistan, cephalosporin resistant isolates belong to the H58 lineage. However, within the H58 lineage, the cephalosporin resistant *S*. Typhi were distributed in different sub-lineages. The Indian isolates belonged to the dominant 4.3.1 lineage II while the isolates from Pakistan clustered with 4.3.1 lineage I. These results are in agreement with previous reports as the MDR associated lineage I is dominant in Pakistan and lineage II (QRDR triple mutants) is more prevalent in India [9, 23–25]. The only exception is the previously reported ceftriaxone resistant *S*. Typhi from Gurgaon, India which originated from lineage 4.3.1.1 [13]. In contrast to the above observations, isolates from Bangladesh, Democratic Republic of Congo (DRC) and Philippines clustered with the non-H58 phylogenetic lineages (26–29).

The evolutionary pattern observed in cephalosporin resistant *S*. Typhi in India may not be driven by acquisition of plasmids already carrying cephalosporin resistance, but by adaptation due to antibiotic usage selection pressure. This is supported by earlier evolutionary events such as the integration of the MDR gene cassette (composite transposon) in the chromosome and the replacement of lineage I by lineage II in India [30]. Further, the emergence and regional dominance of QRDR triple mutant lineage II in India was associated with high fluoroquinolone exposure in the region [6, 24]. Based on available data, it appears that the H58 lineage I with a chromosomal AMR cassette and single QRDR mutation can acquire and maintain extrachromosomal elements, such as plasmids harboring AMR determinants, as in the XDR Pakistan isolates and in the Indian isolate from Gurgaon. Conversely, lineage II strains with double/triple QRDR mutations develop cephalosporin resistance by short term adaptation of resistance plasmids such as IncX3 or IncN. This could probably be the reason for lineage I acquiring IncY to have had potential to cause larger outbreaks and as well its persistence till date, which is in contrast to lineage II in India with lack of potential to spread unlike lineage I in Pakistan. In general, the rise and spread of plasmid mediated AMR in clinical settings is based on the ability of bacteria to compensate for the initial fitness cost imposed by the plasmid [31]. Since the QRDR triple mutant (lineage II) carries the highest fitness cost among fluoroquinolone resistant *S*. Typhi [32], the spread of the plasmid-carrying cephalosporin resistant clone may be limited. Notably, when the lineage can alleviate the cost imposed by the plasmid through compensatory mutations, the spread of these clones will cause even greater therapeutic difficulties.

The global spread of MDR *S*. Typhi was associated with independent acquisitions of IncHI1 plasmids of varying plasmid types (PST) at different locations between the 1970s and 1980s [33]. In contrast, a single plasmid type (PST6) of IncHI1 was responsible for the clonal expansion of H58 haplotype [23, 33]. Similarly in the Pakistan outbreak, a single acquisition event of IncY plasmid harboring *bla*_CTXM-15_ led to the expansion of XDR *S*. Typhi (4.3.1.1P) with a difference of 6 SNPs from the MDR associated *S*. Typhi [9]. Though the fitness of the host strain may have compromised while acquiring the IncY plasmid, the XDR clone appears to have alleviated the cost bycompensatory evolution. As a result, the IncY plasmid appears to be stable and maintained in the genome, as XDR outbreaks are still reported from Pakistan [10]. Our data along with a previous report from northern India [13] indicates that the development of cephalosporin resistance in *S*. Typhi in India has occurred by at least three independent and distinct events. This includes the transfer of IncN plasmid in a single isolate, IncX3 in 4 other isolates and IncY as observed in Pakistan. Plasmid characterization of our study isolates with respect to phylogenetic lineages indicates a short term adaptation of IncX3/IncN plasmids in H58 lineage II (QRDR triple mutant) which are identified sporadically, while successful host adaptation of IncY plasmid in XDR H58 lineage I has led to persistence in Pakistan.

Although our study is limited in sample size and geographical representation, the preliminary observation derived from this study highlights the ability of *S*. Typhi to acquire diverse plasmids to develop cephalosporin resistance. A follow up study on large number of samples, collected evenly across the country, would be required to establish the rate of evolution and clonal expansion in these newly discovered cephalosporin resistant *S*. Typhi.

## Conclusion

Even though ceftriaxone resistant *S*. Typhi are not widely seen in India at present, emergence and spread is possible due to the current high use of azithromycin and ceftriaxone for the treatment of typhoid fever. As a result, there are potential risks for the occurrence of plasmid transmission events. This is based on the following evidence, (i) endemicity of *bla*_SHV_ and *bla_CTX-M-15_* carrying plasmids in *E.coli* and *Klebsiella* sp. in India favoring horizontal gene transfer to *S*. Typhi, (ii) high use of ceftriaxone for the management of complicated typhoid fever posing antibiotic pressure, and (iii) the dominant H58 lineage in India being capable of acquiring plasmid harboring AMR determinants. Considering the prolonged maintenance of newly acquired *bla_CTX-M-15_* carrying IncY plasmid in 4.3.1 lineage I in Pakistan, any trigger could possibly lead to similar events in India. Therefore we propose that MDR H58 lineage II are capable of acquiring MDR plasmids from other Enterobacteriaceae and could potentially cause a large outbreak. Hence, monitoring of cephalosporin resistant *S*. Typhi and its lineages associated with plasmid acquisition is required for early detection, in conjunction with rational use of antibiotics and prevention strategies for control of enteric fever.

## Acknowledgments

This project was supported by grants from Bill and Melinda Gates Foundation, USA (Investment ID INV-009497 OPP1159351) for the Project “National Surveillance System for Enteric Fever in India”

## Author contribution statement

BV conceived and designed the research, JJJ and AKP supervised the research and wrote the manuscript. KV, GK, JJ, VN and AM have reviewed and corrected the manuscript. All authors have read and approved the manuscript.

